# Structural Priming Is Supported By Different Components Of Non-Declarative Memory: Evidence From Priming Across The Lifespan

**DOI:** 10.1101/190355

**Authors:** Evelien Heyselaar, Linda Wheeldon, Katrien Segaert

## Abstract

Structural priming is the tendency to repeat syntactic structure across sentences and can be divided into short-term (prime to immediately following target) and long-term (across an experimental session) components. This study investigates how non-declarative memory could support both the transient, short-term and the persistent, long-term structural priming effects commonly seen in the literature. We propose that these characteristics are supported by different subcomponents of non-declarative memory: Perceptual and conceptual non-declarative memory respectively. Previous studies have suggested that these subcomponents age differently, with only conceptual memory showing age-related decline. By investigating how different components of structural priming vary across the lifespan, we aim to elucidate how non-declarative memory supports two seemingly different components of structural priming. In 167 participants ranging between 20 and 85 years old, we find no change in short-term priming magnitude and performance on perceptual tasks, whereas both long-term priming and conceptual memory vary with age. We suggest therefore that the two seemingly different components of structural priming are supported by different components of non-declarative memory. These findings have important implications for theoretical accounts of structural priming.

---

Structural priming refers to the facilitation of syntactic processing that occurs when a syntactic structure is repeated across consecutive sentences. This can occur for both language comprehension and production; the current article will focus solely on the latter. Structural priming in language production presents behaviourally as an increased tendency to re-use syntactic structures that have been produced either by the speaker or an interlocutor. Such structural persistence has been demonstrated experimentally for different syntactic structures (Bernolet, Collina, & Hartsuiker, 2016; Bock, 1986; Bock & Griffin, 2000), in different languages (Bock, 1986; Hartsuiker & Kolk, 1998; Sung, 2015), and using different priming modalities (Branigan, Pickering, & Cleland, 1999; Branigan, Pickering, Stewart, & McLean, 2000; Hartsuiker, Bernolet, Schoonbaert, Speybroeck, & Vanderelst, 2008). However, although structural priming is a well-established phenomenon, the mechanism underlying this effect is still under much debate (Pickering & Ferreira, 2009).

The debate is largely fuelled by the need for models having to explain not only a single structural priming component, but two: A short-term priming effect and a long-term priming effect (Bock & Griffin, 2000; Chang, Dell, & Bock, 2006; Ferreira & Bock, 2006). The short-term priming effect refers to the priming effect seen for the target sentence immediately following the prime sentence. This is how priming magnitude is most commonly calculated in experimental studies (although see below for exceptions). Earlier studies also reported a “lexical boost” effect, which refers to the increased priming magnitude as occurring due to an overlap of lexical information between prime and target sentences (Pickering & Branigan, 1998). However, a recent study by Bernolet and colleagues (2016) demonstrated that short-term priming effects are observable in experiments with minimal lexical overlap (in this case, the verb was not repeated between prime and target sentences), and occurs for all structures commonly used when studying structural priming (i.e., transitives, datives, and word order for relative clauses). This short-term priming effect, without lexical overlap, is also commonly referred to as abstract priming. In this article, we will focus only on abstract priming by ensuring that the prime and target images do not share a verb or actors.

The second structural priming component, the long-term priming effect, refers to the tendency for participants to increase their use of the primed structure throughout the length of the experimental session, irrespective of the previously presented prime structure (Jaeger & Snider, 2008, 2013; Kaschak, 2007). This suggests that their structural choice is not only influenced by the immediately preceding prime sentence, but also the number of exposures to the primed structure.

The long-term priming effect is commonly explained in terms of a non-declarative, or implicit, learning account. This is motivated by studies showing that participants seem to be unaware both of the priming manipulation and of the fact that they have indeed changed their structural preferences (Bock, 1986; Bock & Griffin, 2000). The current models differ in the mechanisms they propose to explain this non-declarative learning. For example, Chang and colleagues (2006) have proposed an error-based learning account where participants constantly update their predictions based on both prior and recent experience. Jaeger and Snider’s (2013) expectation-adaptation model is similar to this account, except that it does not commit to a specific error-based learning mechanism (although see Jaeger & Snider (2013) for more in-depth comparisons that are beyond the scope of the current paper). However, both accounts find the short-term priming effects more difficult to explain. Jaeger and Snider (2013) suggest that non-declarative memory underlies both the long- and short-term priming effects, but they provide no details about short-term priming even though they do observe a robust effect (i.e., increased probability of re-using the prime structure in the immediately following target response) in experiments 2 and 3. Chang and colleagues (2006) acknowledge that there is “a deep unresolved issue” (pg. 256) and argue that a separate process is required to account for the short-term priming effect. Since then, they have suggested a complementary systems account where short-term priming is due to fast hippocampal learning that interfaces with slow non-declarative learning mechanisms for long-term priming (Chang, Janciauskas, & Fitz, 2012). A recent study by Bernolet and colleagues (2016) tested this theory by measuring both the decay rate of the structural priming effect as well as the decay rate of participants’ explicit recognition of past sentences with the hypothesis that a correlation between the two would support similar underlying mechanisms. Although both showed decay with intervening filler sentences, the strongest priming effect was correlated with the lowest memory scores (specifically, priming for the AUX-PART alternation was stronger that the priming for datives and transitives, while memory for the AUX-PART sentences was worse than for the datives and transitives), providing evidence against this theory. Additionally, an explicit memory theory cannot explain how patients with a deficit in explicit memory still show robust short-term priming effects (Ferreira, Bock, Wilson, & Cohen, 2008; Heyselaar, Segaert, Serge, Kessels, & Hagoort, 2017).

The short-term priming effect has also been explained as residual activation (Malhotra, 2009; Pickering & Branigan, 1998; Reitter, Keller, & Moore, 2011): Recently processed structures remain partially active, increasing the chances of re-use in an upcoming utterance. Although this may explain the short-term priming effect for experiments in which the target immediately follows the prime, there are numerous studies with linguistic or non-linguistic (i.e., time) fillers between prime and target trials that still find a robust “short-term” priming magnitude, even with a week between prime and target (e.g., Bock & Griffin, 2000; Branigan et al., 1999, 2000; Hartsuiker et al., 2008; Kaschak, 2007; Kaschak, Kutta, & Schatschneider, 2011; Reitter, 2008). Malhotra (2009) and Reitter and colleagues (2011) propose an explicit memory basis for this residual activation, which we have argued against above. However, Reitter and colleagues do propose a way in which the short-term residual activation could support long-term non-declarative learning: The spreading residual activation has a power-law decay rate which would predict that this residual activation is never completely lost, allowing a build-up over time with repeated exposures. Hence there is a short-term priming effect due to the previously processed structures still being partially active, which increases the chances of selection when planning the next utterance. A long-term priming effect then is due to the frequency of the structure being “logged” such that repeated retrieval increases the base activation of a structure, so that more frequent structures have a higher base activation, and hence a higher chance of selection in an upcoming utterance.

In this study, we propose a merge between different aspects of the above models but one which is exclusively based in non-declarative memory. For short-term priming we propose a residual activation account based in non-declarative memory (similar to that proposed by (Malhotra, 2009; Pickering & Branigan, 1998; Reitter et al., 2011) and for long-term priming a non-declarative learning account (similar to that proposal by Chang et al., 2006; Jaeger & Snider, 2013). The information transfer between these components is based on the information transfer proposed above in the Reitter model. The key difference with our account of structural priming is that we base everything in non-declarative memory. This proposal is not new, and has been explained in depth in MacDonald (2013) for general language processing. In the article, the author refers to Easy First (short-term) and Plan Reuse (long-term) mechanisms and provides examples not only for general language production but also for other cognitive behaviours, such as motor planning. Indeed, it seems logical to apply these mechanisms to structural priming given its characteristics, and yet no study, to our knowledge, has tested this empirically. We will next explain why we believe structural priming is based solely in non-declarative memory.

Implicit, non-declarative memory has been defined as the unconscious memory of events that participants may not consciously recollect (Graf & Schacter, 1985; Schacter & Tulving, 1994). This is tested indirectly by having participants perform a task in which no apparent reference is made to any prior episode. For example, the word-stem completion task consists of three letter word-stems that the participant is asked to complete with the first word that comes to mind. However, unbeknownst to the participant, these stems can all be completed using words the participant has been exposed to earlier, via a questionnaire or other seemingly unrelated task within the same study session. Tasks can also test more complex relationships. For example, in serial reaction time tasks participants think they are completing a reaction time task (responding to a stimulus on the screen as fast as possible) but in fact the stimuli presented have an underlying pattern that the participant unconsciously learns. This learning results in decreased reaction times over the length of the session as the participants are able to unconsciously predict the upcoming stimuli. Non-declarative memory performance is therefore measured as an increased efficiency (i.e., increased accuracy or decreased latency) in processing information that participants have been exposed to at an earlier stage, and is attributed to slow-decaying residual activation. This type of memory has also been referred to as procedural memory (Cohen & Eichenbaum, 1993).

Studies in the memory literature have suggested that non-declarative memory is made up of (at least) two components (Gabrieli, 1998; Gupta & Cohen, 2002; Squire, 2004). *Conceptual memory* (also referred to as skill learning) supports the learning of statistical covariations and dependencies between stimuli (e.g. serial reaction time tasks), whereas *perceptual memory* (also referred to as repetition priming) maintains residual activation of a recently processed stimulus. Short-term structural priming is proposed to work in a similar way: The residual activation of a previously processed syntactic structure that persists, represented as an increased chance in reusing that structure in the following utterance. Tasks designed to investigate perceptual memory measure how previous exposure to a specific stimulus (e.g., a word) facilitates later processing of that word or a related item (e.g. word-stem completion and fragmented identification tasks). Based on the definitions of these components, we propose that long-term priming is most likely supported by conceptual memory, whereas short-term priming most likely supported by perceptual memory. We propose that non-declarative memory supports both temporal characteristics of structural priming, yet different components of this memory system underlie the different components of structural priming. To further support our proposal, we turn to how the memory system changes as we age.

It is well established in the literature that declarative memory declines with age. This decrease in the ability to encode and retrieve explicit information has been linked to decreases in hippocampal (Golomb et al., 1993; although see Raz et al., 2003) and medial temporal lobe volume (Bailey et al., 2013) as well as impaired functioning of the right frontal regions (Stuss, Craik, Sayer, Franchi, & Alexander, 1996). For non-declarative memory, for quite some time there was a consensus that this system was not susceptible to age related decline. However, together with the discovery that there are subsystems within non-declarative memory, evidence has emerged that different neural networks support these systems, and that the systems could therefore be differentially susceptible to age-related decline. Neuroimaging studies of healthy older adults and patient studies have shown that conceptual and perceptual memory have distinct neural correlates. Perceptual memory is associated with activity in the posterior cortical regions (Bäckman, Almkvist, Nyberg, & Andersson, 2000; Squire et al., 1992), whereas conceptual memory is associated with a subcortical-cortical network in which the striatum is a central component (Lieberman, Chang, Chiao, Bookheimer, & Knowlton, 2004). There are studies showing age-related decline in the striatum (Bäckman, Nyberg, Lindenberger, Li, & Farde, 2006; Raz et al., 2003), which would affect conceptual but not perceptual memory. However, a review of behavioural studies looking at the aging effect of conceptual and/or perceptual studies draws equivocal conclusions. This is mainly due to different experimental designs: Most aging studies take a younger age group (on average 25 years) and compares it directly to an older age group, which can vary from early 60’s (Howard, Heisey, & Shaw, 1986; Neger, Rietveld, & Janse, 2014; Schugens, Daum, Spindler, & Birbaumer, 2007) to late 80’s (Davidson, Zacks, & Ferreira, 2003; Davis et al., 1990; Karlsson, Adolfsson, Börjesson, & Nilsson, 2003; Light, Kennison, & Healy, 2002; Light, La Voie, Valencia-Laver, Albertson Owens, & Mead, 1992). Especially at the older age group, it is unclear at what point non-declarative memory may start to decline, which may explain why some find a significant difference with the younger age group and others do not. To our knowledge, only one study (Maki, Zonderman, & Weingartner, 1999) tested participants from each decade from 20’s until 80’s in a perceptual memory task (Fragmented Object Identification) and a conceptual memory task (Category Exemplar), and conclude that perceptual memory declines with age, whereas conceptual memory stays intact. This is in stark contrast to our hypothesis and the neuroimaging data described above, however, as it is one study in a field of methodologically inconsistent studies we propose to measure the perceptual and conceptual memory of our participants, in order to make a more direct link to their structural priming ability.

The aim of the current study is thus to test our hypothesis that non-declarative memory supports both key temporal characteristics of structural persistence. In contrast to current models of structural priming (Chang et al., 2006; Jaeger & Snider, 2013; Pickering & Branigan, 1998), we propose that different subcomponents of non-declarative memory underlie both short-term and long-term structural priming. Specifically, we propose that perceptual memory underlies short-term priming, while conceptual memory underlies long-term priming. In the current study we therefore tested structural priming in 167 participants aged between 20 and 85 years. If our hypothesis about the role of conceptual and perceptual memory is accurate, we should observe that, as the age of the participant increases, their long-term priming magnitude declines whereas their short-term priming magnitude remains unaffected. Reviewing the structural priming literature does not lead to any clear conclusions on how priming ability changes with age: The studies that show intact priming in older adults (Ferreira et al., 2008; Hardy, Messenger, & Maylor, 2017; Hardy, Wheeldon, & Segaert, 2018; Hartsuiker et al., 2008) are in contrast with studies showing a decline in priming ability in older adults (Heyselaar et al., 2017). Additionally, these studies have focused on short-term priming, with little to no results indicating how long-term priming may change with age.

We also measured the participants’ performance on well-established memory tests designed to measure conceptual and perceptual memory. The tasks included in our non-declarative memory battery have been frequently used in the literature: A word-stem completion task (Light & Albertson, 1989; Light & Singh, 1987), a fragmented identification task (Au et al., 1995; Mitchell, 1989), and a serial reaction time task (Nissen & Bullemer, 1987). The serial reaction time task measures conceptual memory. We included not one but two perceptual memory tasks: The word-stem completion task and the fragmented identification task. The former is a prominent task in the literature, however, a meta-analysis (La Voie & Light, 1994) has suggested that the word-stem completion task is prone to declarative memory contamination.

In line with neuroimaging data, we predict that participants will show a decline in the conceptual task (serial reaction time task) and show no decline in the perceptual tasks (word-stem completion and fragmented identification tasks). A demonstration of a comparable effect of aging on these tasks and on the effects of long- and short-term structural priming will provide evidence in support of our proposal that different components of non-declarative memory underlie these aspects of structural persistence. Additionally, our study will be one of the first to show different effects of age on the two temporal characteristics of structural priming, an important discovery that needs to be included in future models of structural priming.

## Materials And Methods

### Participants

167 participants (62 men) were recruited through the Patient and Lifespan Cognition participant database of the School of Psychology at the University of Birmingham and through flyers and advertisements in and around the University of Birmingham. Most participants were tested at the university and some were tested at their place of work or in their homes. We attempted to obtain an equal number of participants for each decade of life: 20 – 29 years (*n* = 25), 30 – 39 years (*n* = 24), 40 – 49 years (*n* = 27), 50 – 59 years (*n* = 23), 60 – 69 years (*n* = 31), and 70 – 85 years (*n* = 37). All participants were required to have British English as their mother tongue and have at least a university degree in order to minimize education-related differences in performance. At the time of testing, no participants reported any neurological deficits or psychiatric disorders. The study was approved by the research ethics board of the University of Birmingham. All participants were paid for their participation.

### Study Design & Apparatus

All participants completed one structural priming task, three non-declarative memory tasks (word-stem completion, fragmented identification task, and serial reaction time task), one declarative memory task, two verbal working memory tasks (backward digit span and subtract-2 span task), and one verbal IQ task (national adult reading task) in one 1.5-hour session. Figure 1 illustrates the order of events. All participants completed the tasks in the same order.

**Figure 1.**
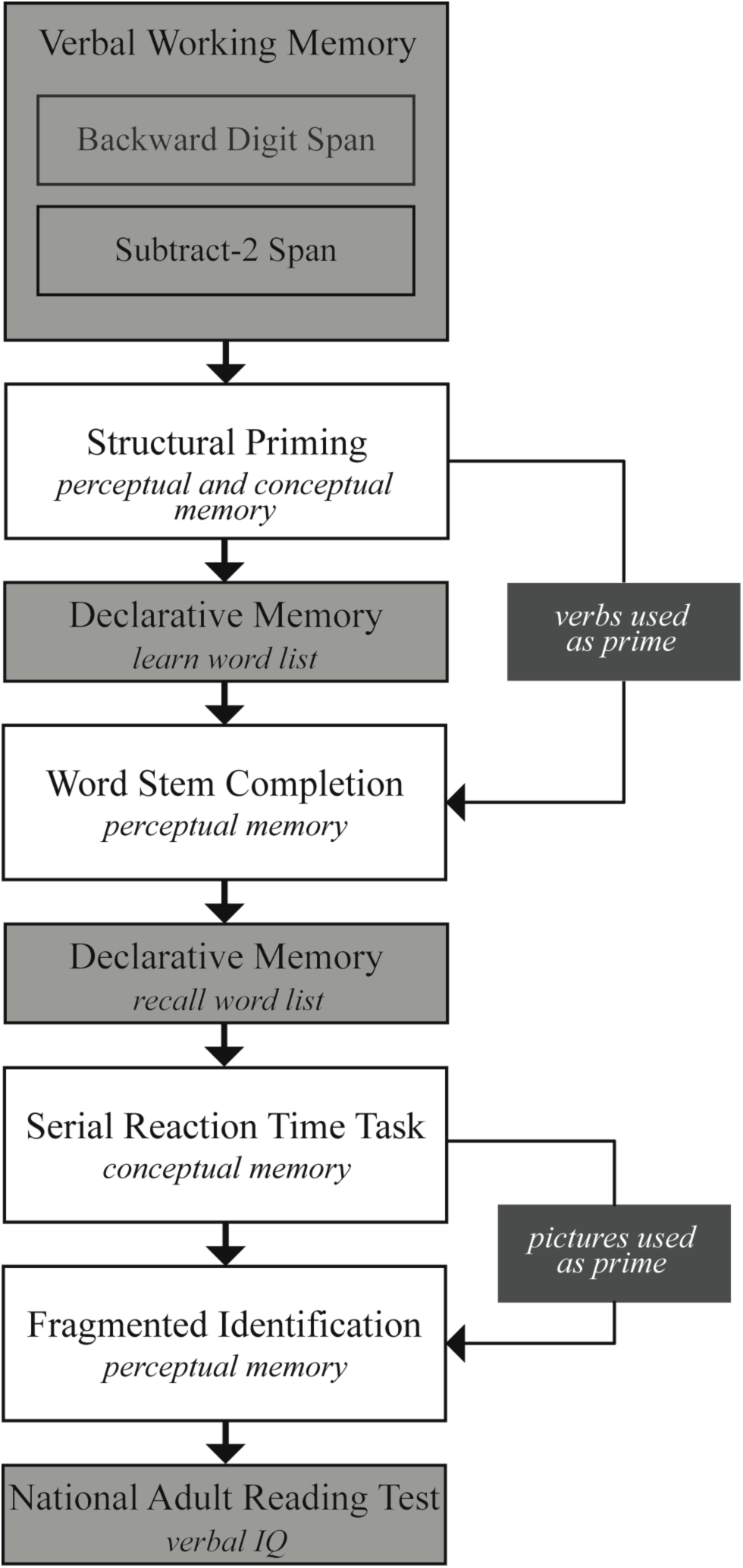
All participants completed all tasks in the order illustrated here. Grey boxes represent control tasks, measuring working memory, explicit memory, and verbal IQ. White boxes represent the non-declarative memory tasks. Some tasks, in addition to measuring non-declarative memory, also acted as primes for future non-declarative memory tasks (black boxes).

All tasks were completed on a Dell Latitude E5470 Laptop (14’’ screen) using E-Prime (Schneider, Eschman, & Zuccolotto, 2002).

We included control measures of declarative memory, verbal IQ, and working memory and included each participant’s score in our statistical models to partial out their contribution to performance in our non-declarative memory tasks.

### Structural Priming Task

This task was based on a task by Menenti, Gierhan, Segaert, and Hagoort (2011).

The pictures used in this task were taken from Segaert, Menenti, Weber, and Hagoort (2011). The stimulus pictures depicted 40 transitive events such as *to chase, to interview* or *to serve* with a depiction of the agent and patient of this action. Transitive pictures were used to elicit transitive prime and target sentences. Each transitive picture had three versions: one grayscale version and two colour-coded versions with a green and a red actor. Participants were instructed to describe pictures with one sentence, naming the green actor before the red actor if the actors were depicted in colour. This allowed us to manipulate, for colour-coded primes, whether the prime sentence produced had an active (e.g., *the man kisses the woman*) or a passive (e.g., *the woman is kissed by the man*) syntactic structure.

Each transitive event was depicted by two pairs of adults and two pairs of children. One male and one female actor were shown in each picture, and each event was depicted with each of the two actors serving as the agent. To prevent participants forming strategies, the position of the agent (left or right) was randomized. Fillers elicited intransitive sentences, depicting events such as *running, singing,* or *bowing* with one actor (in greyscale or green).

Each experimental trial consisted of a prime (a coloured picture) followed by a target (a greyscale picture). There were 20 passive prime trials (a passive picture followed by a transitive greyscale target), 20 active prime trials (an active picture followed by a transitive greyscale target), and 20 baseline trials (an intransitive picture followed by a transitive greyscale target), all randomized in one experimental session. The baseline trials allowed us to measure the frequency of producing active and passive transitives on subsequent targets without any immediate prior influence. We also included 20 filler trials (an intransitive picture followed by an intransitive greyscale target). In total, therefore, we had 80 trials consisting of 100 transitive pictures and 60 intransitive pictures.

The timing of each prime, target or filler trial was as follows: Participants were initially presented with a neutral verb (to be used in an upcoming utterance; e.g. “to run”, “to chase”, etc.) for 500ms. After 500ms of black screen a coloured picture would appear. Participants were instructed to describe the picture following the rules described above. The picture was presented until the participant responded (with a time-out after 12 seconds). There was an intertrial interval of 1500 - 2000ms (jittered) before the next verb was presented. Coloured and greyscale pictures were alternated until all pictures were described. The task took a total of 15 minutes to complete.

Responses were manually coded by the experimenter as either active or passive. Trials in which the descriptions did not match one of the coded structures were discarded (6.96% of the data). Target responses were included in the analysis only if 1) both actors and the verb were named (a sentence containing only one of the actors does not qualify as a transitive sentence) and 2) the structures used were active or passive.

The proportion of passives produced after a passive prime compared to baseline (measurement of short-term priming), as well as the number of passives produced throughout the length of the task regardless of prime type (measurement of long-term priming) were taken as variables of interest.

### Non-declarative Memory Tasks

We will describe the tasks in the order they were presented in the experiment. We included two perceptual memory tasks: The word-stem completion task (verbal memory) and the fragmented identification task (visual memory), and one conceptual memory task: The serial reaction time task.

#### Word-stem completion (WSC) task

This task was developed to test the non-declarative memory for words. This task, including the number of target word-stems used, is based on those described in Davis and colleagues (1990) and Fleischman, Wilson, Gabrieli, Bienias, and Bennett (2004) but adapted for use on a computer.

Participants were presented with 20 three-letter word-stems on the computer screen and instructed to complete the words by typing in their answer using the keyboard. Participants were encouraged to use the first word that came to mind.

10 of the word-stems could be completed using the verbs used in the structural priming task (randomly selected from the list of verbs each specific participant used) and 10 using novel words (randomly picked from a list of 33 stems). The 33 word-stems to be completed by novel words were selected from a word-stem database by Migo, Roper, Montaldi, and Mayes (2010). The novel word-stems could not be completed with any of the words the participants had been exposed to until this point in the session. The word frequency of the most common completions for the test stems and novel stems did not significantly differ (Independent samples *t*-test, t(42) = -.874, *p* =.300).

The 20 three-letter word-stems were presented one at a time, in a random order, and were only replaced with a new stem once the participant entered their completed word. The task took 5 minutes to complete.

The number of word-stems completed with verbs primed in the structural priming task as well as the reaction times were taken as variables of interest. Reaction times were trimmed to 2.5 standard deviations for each age group (11 out of 352 data points were removed; 3.13% of the data).

#### Serial reaction time (SRT) task

This task was developed to test statistical co-occurrences of temporally separated stimuli (Nissen & Bullemer, 1987) and therefore is a measure of conceptual memory. This task is based on a task described by Neger, Rietveld, and Janse (2014).

Participants were presented with a 3 x 3 grid on the computer screen that was filled with the digits 1 to 9. A picture would be presented on one of the 9 locations, and participants were instructed to press the corresponding number key as fast as possible. Participants were instructed to respond using the number pad, such that the keys on the number pad correspond to the same spatially located key on the grid. Crucially, the location of the subsequent picture could be predicted based on the location of the current picture. The pictures used were not relevant for this task, however, for each participant the same picture would appear at the same location for the duration of the task. Pictures and their locations were randomized between participants.

Similar to Neger and colleagues (2014), the task was composed of blocks and split into an exposure phase, a test phase, and a recovery phase. During the exposure phase, participants could learn the underlying pattern by picking up on the co-occurrence probabilities of the locations. In total, the exposure phase consisted of 16 predictable blocks. Within each block, all location combinations were repeated once, resulting in 128 exposure trials (8 x 16). The test phase consisted of two unpredictable blocks, resulting in 16 test trials (8 x 2). In these unpredictable trials, a new underlying pattern was used. Participants who implicitly learnt the patterns in the exposure phase should show a drop in performance as they would need to correct their predictions during the test phase, resulting in a slowed response to the second picture. This measure of learning is widely accepted in the literature on conceptual memory (Janacsek & Nemeth, 2013).

A picture would only appear on the specific location 500ms after the onset of the visual display and the task only proceeded if the participant pressed the appropriate target number. The new location would only be revealed 500ms after the previous picture disappeared, a time interval that has previously been shown to be necessary to successfully allow prediction effects in older adults (Salthouse, McGuthry, & Hambrick, 1999). A fixation cross appeared for 2500ms between blocks for all phases. In total this task took 20 minutes to complete.

To assess skill learning, we measured latencies from the picture presentation to the subsequent correct button press. To correct for any changes in response time due to age, we calculated facilitation scores for each participant. Facilitation score was calculated by dividing the RT to a location by the RT to the subsequent location. Facilitation scores were trimmed to 2.5 standard deviations for each age group (5 out of 176 data points were removed; 2.84% of the data).

As each location primed the subsequent location, a single location acted as both a target on the current trial and as a prime for the next (contrary to previous versions of this task where a location can be only a prime or a target, not both). To measure skill learning, we focused our analysis only on locations where the prime location was unpredictable (<25%) and the targets were highly predictable (>75%). This decision was reached after data-collection to minimize the noisiness of the data.

We calculated percent change for each participant’s facilitation score between phases and entered this as the dependent variable in the model.

#### Fragmented identification task

This task was developed to test the non-declarative memory for pictures. This task is based on a task by Kessels, Remmerswaal, and Wilson (2011) but adapted for use on a computer.

Participants were shown a set of 16 line drawings, in a sequence of 8 pictures of decreasing degradation. The participant had been previously exposed to 8 of the line drawings during the SRT task while the remaining 8 were novel line drawings the participant had not seen before. The dependent measure was the level of degradation at which accurate identification was possible. The pictures were selected from the bank of standardized stimuli (BOSS) picture database (Brodeur, Dionne-Dostie, Montreuil, & Lepage, 2010; Brodeur, Guérard, & Bouras, 2014). We used naming frequency from the BOSS database to pick 18 pictures that were 1) named with only one name and 2) named using the same name 100% of the time. This was to ensure we used pictures with the highest rate of identification. All pictures had comparable complexity scores (M: 2.29; SD: 0.402).

Fragmentation of the pictures was done manually following the methods described by Snodgrass, Smith, and Feenan (1987) but briefly: Pictures were placed into a 16 x 16 block grid. The locations of blocks that contain picture information were then identified. Blocks with picture information were randomly selected to be erased to produce eight levels of fragmentation per picture. Level 8 is the complete picture and Level 1 is the most fragmented picture, containing only 8% of the original picture. Contours of the picture were fragmented separately to ensure that the outline of each picture was also fragmented to the same extent as the rest of the picture, i.e., at Level 1, only 8% of the contours were visible. This prevented the participants from being able to identify a picture at a low fragmentation purely because the entire outline of the picture was complete.

The participant was instructed to type the name of the picture. If the answer was incorrect, the participant would be shown the next picture in the fragmented sequence. If the participant was correct, they would be moved on to the next novel object to identify. Each picture in the sequence was presented for at least 3 seconds and until the participant indicated they were ready to see the next picture in the sequence (hence a response was not required for each fragment in the sequence). In total the task took 10 minutes to complete.

The dependent variable in this task was therefore the sequence number at which the participant correctly identified the picture, for each line drawing.

### Additional Measures

#### Verbal working memory tasks

Waters and Caplan (2003) reported that test-retest reliability is considerably better when performance across several verbal working memory tasks are averaged to yield a composite span score. We chose to have participants complete the backward span and subtract-2 span tasks and use their composite score in further analysis. Waters and Caplan (2003) illustrated how these two tasks have the highest correlation of the seven verbal working memory span tasks tested (Pearson’s r = .71, *p* < .05). The tasks used here are based on those described in Waters and Caplan (2003) but adapted for a computer.

##### Backward digit span task

In this task, on each trial, the participants were required to repeat a series of digits in reverse order of presentation. The stimuli were digits drawn from the digits 1 to 9 and presented randomly.

##### Subtract-2 span task

In this task, on each trial, the participants were required to repeat a series of digits after subtracting 2 from each digit. The stimuli were digits drawn from the digits 2 to 11 and presented randomly.

Participants were tested on span sizes 2 to 8 in each of the two verbal working memory tasks. For both tasks, there were five trials at each span size. The participants were required to repeat all the items in the trial in the correct serial order to obtain credit for the trial. They were instructed to submit a blank answer if they could not remember what the item was. The items were presented at the rate of one per second. In total the verbal working memory tasks took 15 minutes to complete.

Span in both tasks was defined as the longest list length for which the participants correctly recalled all the items in the correct serial order on three out of the five trials. An additional 0.5 was added if the participants were correct on two out of the five trials at the next span size. The dependent variable is thus the average span of the two verbal working memory tasks.

#### Declarative memory task

The declarative memory task is based on one used in the Weschler Memory Scale (WMS-III; Wechsler, 1997), but adapted for computer use. Before the WSC task started, participants were presented with a list of 10 words and explicitly told to remember these words. After the participants completed the WSC task, they were asked to type out as many of the to-be-remembered-words as possible. The number of correctly remembered words was used as a measure of declarative memory.

#### Verbal IQ

Verbal IQ was measured using the National Adult Reading Test (NART). This task is based on Nelson and Willison (1991). The participant is given a list of 50 words and asked to read them aloud. Performance is based on whether each word was pronounced correctly. Marking for the NART was done offline by the same experimenter (E.H.) for all participants.

### Data Analysis

All of the data analysis was done coding age as a linear variable, not binned in the age groups described under *Participants*. Therefore, *Age* (factor) or age (in text) refers to age as a linear variable. *Age Range* (factor) or age group (in text) refers to age binned per decade.

#### WSC and Fragmented Identification

These two non-declarative memory tasks were analysed using mixed-effects models, using the lme4 package (version 1.1-10; Bates, Maechler, & Bolker, 2012) in R (R Core Development Team, 2011). We used a maximal random-effects structure as was justified by the data (Barr, Levy, Scheepers, & Tily, 2013): the repeated-measures nature of the data was modelled by including a per-participant and per-item random adjustment to the fixed intercept (“random intercept”). We attempted to include as many per-participant and per-item random adjustments to the fixed effects (“random slopes”) that converged and did not cause overfitting errors. We began with a full model and then performed a step-wise “best-path” reduction procedure, removing interactions before main effects, to locate the simplest model that did not differ significantly from the full model in terms of variance explained (as described in Weatherholtz, Campbell-kibler, and Jaeger, 2013) using the drop1 function from the stats package (version 3.4.2). For the Fragmented Identification task, we used a Poisson model to better model the count nature of the dependent variable.

#### SRT Task

This task was analysed using linear models, as there were no repeated measures once percentage change was calculated, using the stats package in R. We began with a full model and then performed a step-wise “best-path” reduction procedure similar to the one described above. P values were obtained using the Anova function from the car package (version 2.1-1; Fox & Weisberg, 2011) using Wald Chi-Square tests (Type III).

#### Structural Priming

This task was analysed using a generalised additive mixed model (GAMM), using the mgcv package (version 1.8-22; Wood, 2017) as previous experiments have shown that long-term priming mirrors a growth-curve more than a linear correlation over the length of the experimental session (Heyselaar, Hagoort, & Segaert, 2015; Segaert, Wheeldon, & Hagoort, 2016). We used a binomial response variable to code *Target* (active (0) or passive (1)). The gam.check function was used to run diagnostic tests on the subsequent models. Model comparisons were done using the anova function from the stats package (version 3.4.2).

## Results

### General Descriptives

Table 1 shows the average score on each of the three control measures for each age group included in the analysis. Figure 2 shows the variation of each of the three control measures with age.

**Figure 2.**
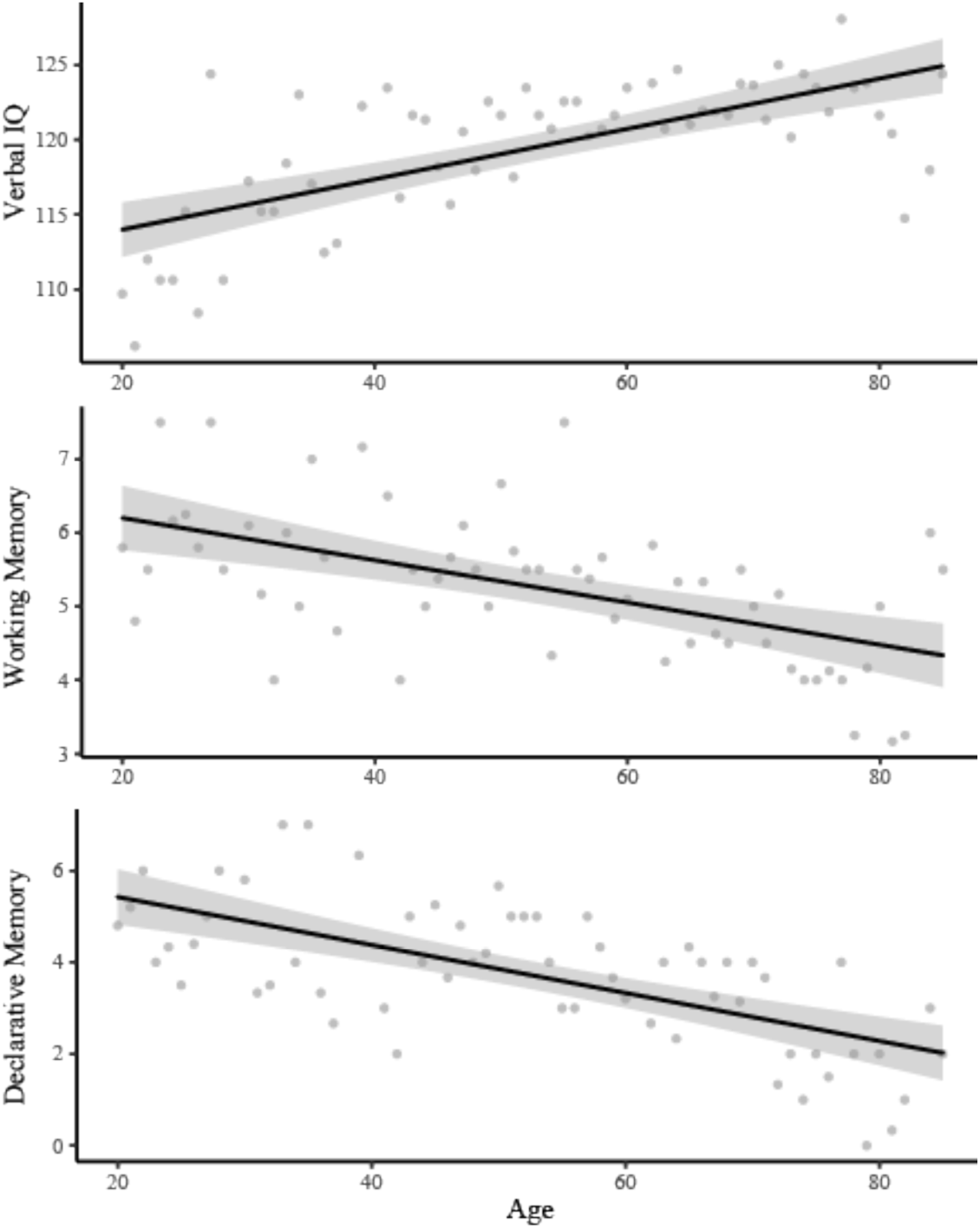
Results of the three control measures. The measure of verbal IQ (NART) showed a significant increase with age whereas the working memory and declarative memory tasks showed significant decreases with age. Error clouds represent standard error.

**Table 1.** Performance of each age group in each of the three control measures.

| Age Range | N | Verbal IQ | Working Memory | Declarative Memory |
| --- | --- | --- | --- | --- |
| 20 – 29 | 25 | 110.14 (4.75) | 5.78 (1.16) | 4.76 (1.79) |
| 30 – 39 | 24 | 116.90 (5.13) | 5.64 (1.34) | 4.66 (2.33) |
| 40 – 49 | 27 | 119.47 (4.62) | 5.46 (1.15) | 4.22 (2.12) |
| 50 – 59 | 23 | 121.01 (3.13) | 5.52 (1.11) | 4.43 (1.75) |
| 60 – 69 | 31 | 122.86 (3.44) | 5.11 (0.89) | 3.32 (1.92) |
| 70 – 85 | 37 | 122.12 (4.18) | 4.27 (1.11) | 2.00 (1.72) |

*Note.* Value represents the mean for each age group, with the standard deviation in parenthesis.

We observe a significant decrease in verbal working memory (Pearson’s r = -0.39, *p* < .001) and declarative memory (Pearson’s r = -0.45, *p* < .001) with age, which is consistent with the literature (Fleischman & Gabrieli, 1998; Fleischman et al., 2004). We observe a significant increase in verbal IQ (Pearson’s r = 0.63, *p* < .001) with age, consistent with findings of increased vocabulary size with age (Brysbaert, Stevens, Mandera, & Keuleers, 2016).

We will describe the results in the non-declarative memory tasks by component. We will focus on the perceptual memory tasks first, then the task that measures conceptual memory, and finally the structural priming task to determine whether the long-term and short-term priming magnitudes match the age-related patterns seen in the non-declarative memory tasks.

### Perceptual Memory - Word-stem Completion (WSC)

There was no significant effect of age on the number of word-stems participants completed with words primed in the structural priming task (Pearson’s r = -0.10, *p* = .076), although there was a trend that older participants completed less word-stems. Table 2 provides the summaries for each age group, although the analysis itself was done on age as a linear variable.

**Table 2.** Number of word stems completed with primed words by each age group. The table lists the average number of word-stems completed with the primed word for each age group. Highest obtainable score is 10.

| Age Range | Average | Standard Deviation |
| --- | --- | --- |
| 20 – 29 | 3.77 | 1.74 |
| 30 – 39 | 3.76 | 1.83 |
| 40 – 49 | 4.21 | 1.50 |
| 50 – 59 | 3.98 | 1.70 |
| 60 – 69 | 3.66 | 1.66 |
| 70 – 85 | 3.35 | 1.35 |

The internal consistency of the response time data was good (Cronbach’s ɑ: 0.95). We used a linear mixed effects model to analyze the response time data. The full model contained two-way interactions of *Primed* (primed vs. not primed word-stems; sum-contrast coded) with *Age* and each of the three control measures as well as two-way interactions of *Age* with each of the three control measures. Interaction effects between *Primed* and *Age* crucially capture the age-related effects on perceptual memory for words. We included no random slopes as then the model would not converge. Table 3 shows the results of the best model, which included main effects of *Primed, Age, Working Memory* (composite score), and *Declarative Memory* (# of words correctly recalled), and a two-way interaction of *Age* and *Working Memory* and a two-way interaction of *Primed* and *Verbal IQ*.

**Table 3.** Summary of the best linear mixed effects model for the Word-Stem Completion Task, modelling the response time to word-stem completion.

|  | coefficient | SE | df | t value | p value |  |
| --- | --- | --- | --- | --- | --- | --- |
| Intercept | 1581.28 | 306.16 | 158.10 | 5.17 | < .001 | *** |
| Primed | -414.64 | 41.00 | 159.46 | -10.11 | < .001 | *** |
| Verbal IQ | -20.64 | 15.63 | 157.31 | -1.32 | .189 |  |
| Age | 26.57 | 5.76 | 157.42 | 4.61 | <.001 | *** |
| Working Memory | 346.91 | 165.55 | 161.12 | 2.10 | .038 | * |
| Declarative Memory | -56.30 | 34.89 | 158.20 | -1.61 | .109 |  |
| Primed * Verbal IQ | 13.15 | 6.87 | 159.23 | 1.92 | .057 |  |
| Age * Working Memory | -9.74 | 3.14 | 163.12 | -3.10 | .002 | ** |
| N = 322 *** < .001 ** < .01 * < .05 |  |  |  |  |  |  |

There is a significant effect of *Primed* on response time such that participants were faster to complete the word-stem with a primed word than with a novel word (Figure 3). The main effect of *Age* was due to a slower response time as the participants increased in age, regardless of whether they completed the word-stem with a primed or not primed word. This could be due to decreased processing speed, as is established for older participants (Salthouse, 1996), or due to the decreased familiarity of typing on a keyboard for older participants, which would affect these participants equally in both conditions. The interaction with *Working Memory* illustrates a similar concept: Participants with a higher working memory composite score were faster at completing the stems (regardless if primed or not), and this interacted with *Age* as working memory capacity decreases with age (as shown in Figure 2).

**Figure 3.**
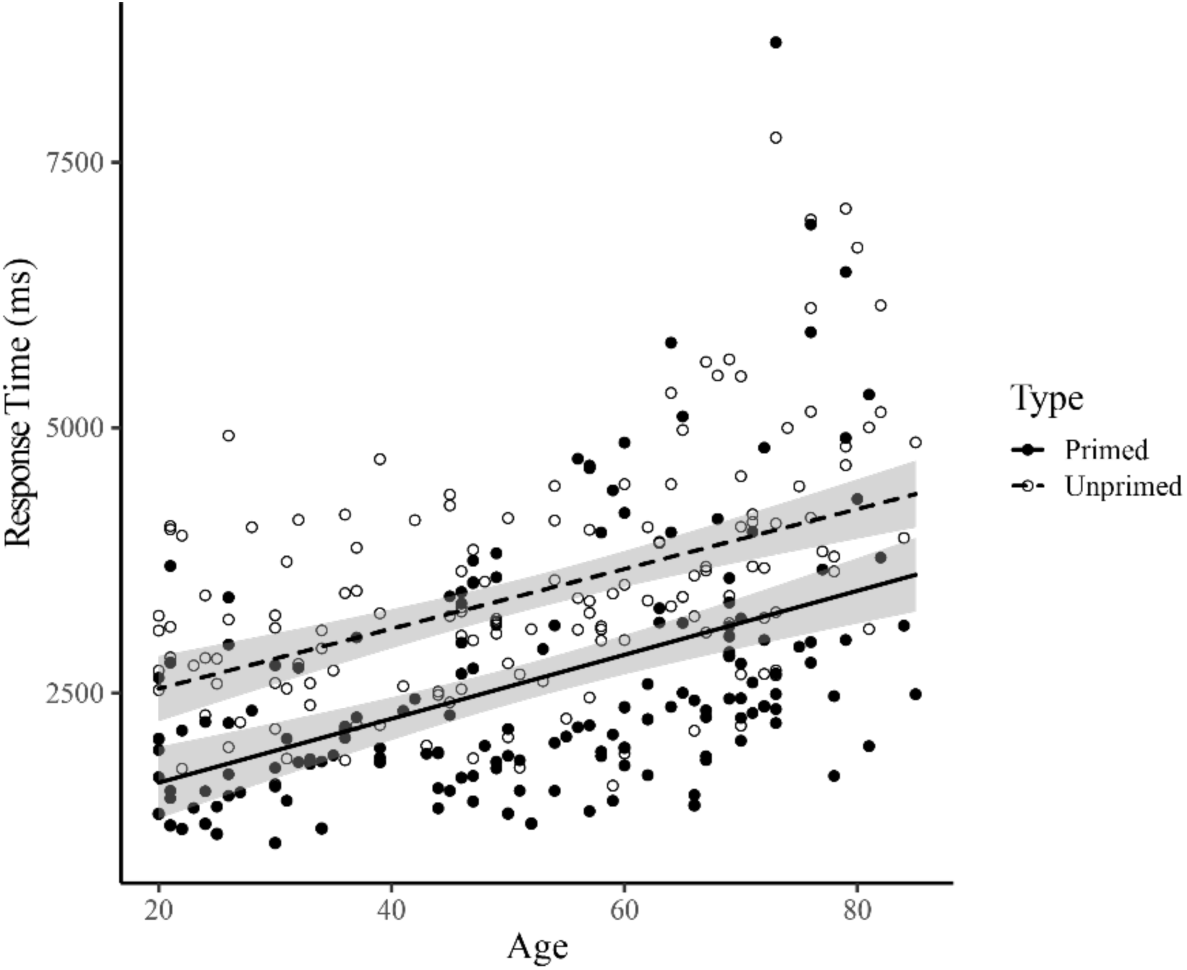
Response Time for the Word-stem Completion Task. Participants were increasingly slower with increasing age (*p* < .001), however, they were consistently faster at completing primed words compared to not primed words (*p* < .001) regardless of age. There was therefore no effect of age on perceptual memory performance in this task. Error clouds represent standard error.

Overall, we do not see an effect of age on perceptual memory performance in this task and we do not see strong evidence in favour of declarative memory contamination for the prime effects (*p* = .097).

### Perceptual Memory - Fragmented Identification

We used a Poisson regression (also known as a log-linear model) to analyse the data, as this better captures count data. The dependent variable was the number of pictures the participant needed to see before they correctly identified it. The internal consistency of the data was good (Cronbach’s ɑ: 0.99). The full model contained a two-way interaction of *Primed* (not primed vs. primed pictures; sum-contrast coded) and *Age.* We also included two-way interactions between *Age* and each of the three control measures. We included no random slopes as we received errors of overfitting. We included a random intercept for items. Table 4 shows the results of the full model, which was also the best model in terms of variance explained.

**Table 4.**
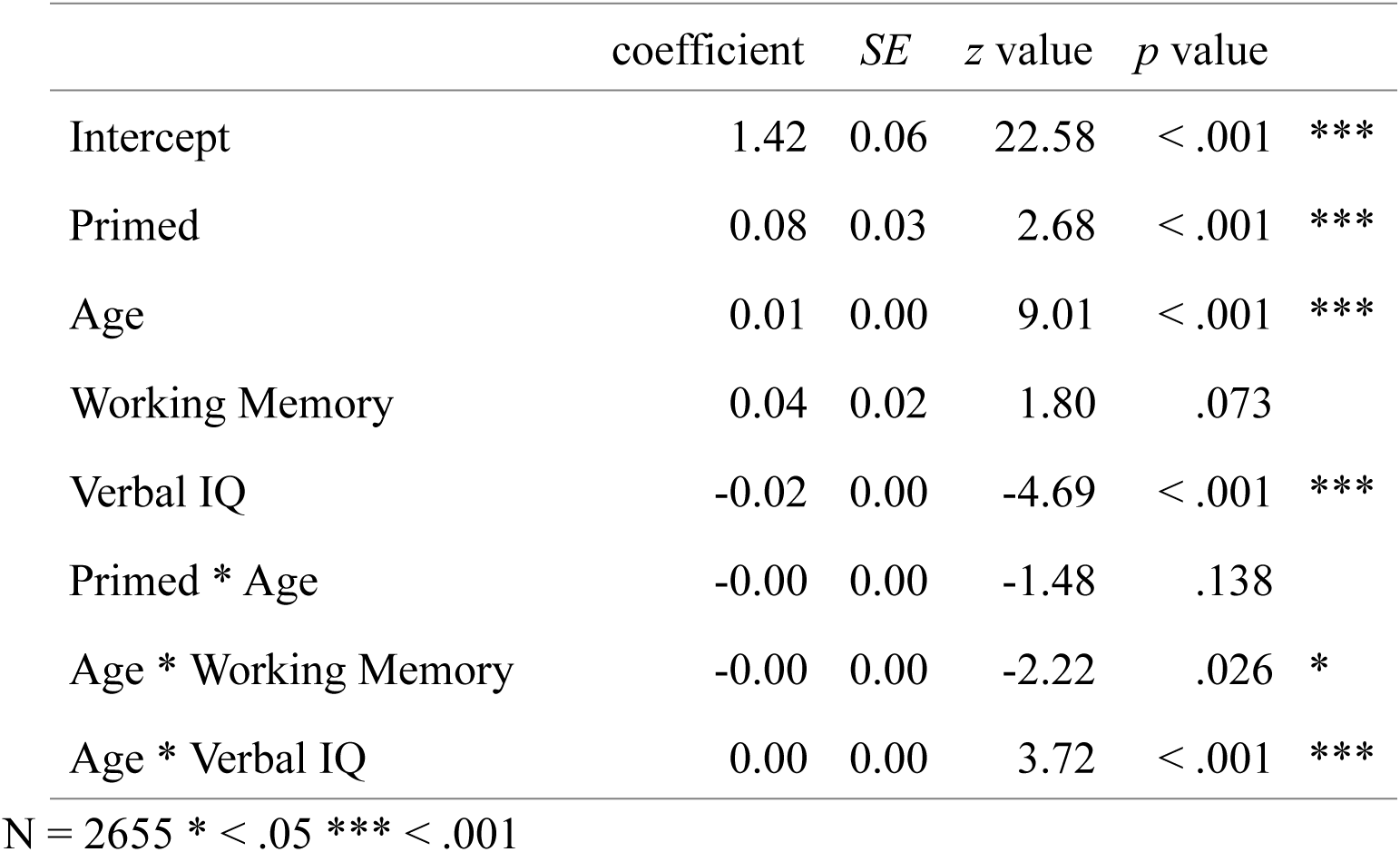
Summary of the best linear mixed effects model for the Fragmented Identification Task. The number of pictures the participant needed to see before they were able to correctly identify the picture is the dependent variable in this model.

|  | coefficient | SE | z value | p value |  |
| --- | --- | --- | --- | --- | --- |
| Intercept | 1.42 | 0.06 | 22.58 | < .001 | *** |
| Primed | 0.08 | 0.03 | 2.68 | < .001 | *** |
| Age | 0.01 | 0.00 | 9.01 | < .001 | *** |
| Working Memory | 0.04 | 0.02 | 1.80 | .073 |  |
| Verbal IQ | -0.02 | 0.00 | -4.69 | < .001 | *** |
| Primed * Age | -0.00 | 0.00 | -1.48 | .138 |  |
| Age * Working Memory | -0.00 | 0.00 | -2.22 | .026 | * |
| Age * Verbal IQ | 0.00 | 0.00 | 3.72 | < .001 | *** |

N = 2655 \* < .05 \*\*\* < .001

The model shows a main effect of *Primed* such that more of the not primed than the primed pictures need to be seen before the picture was correctly identified (Figure 4). There is also a main effect of age: More of the picture needed to be seen as the participants increased in age, regardless of whether the picture was primed or not. The interaction with *Verbal IQ* illustrates that participants with a higher verbal IQ needed to see more of the picture in order to identify it (regardless if primed or not), and this interacted with *Age* as verbal IQ increases with age (as shown in Figure 2). The interaction of *Age* and *Working Memory* is mainly driven by the fact that 60 – 69-year olds with better working memory need to see less of the picture (regardless if the picture was primed or not) in order to correctly identify it.

**Figure 4.**
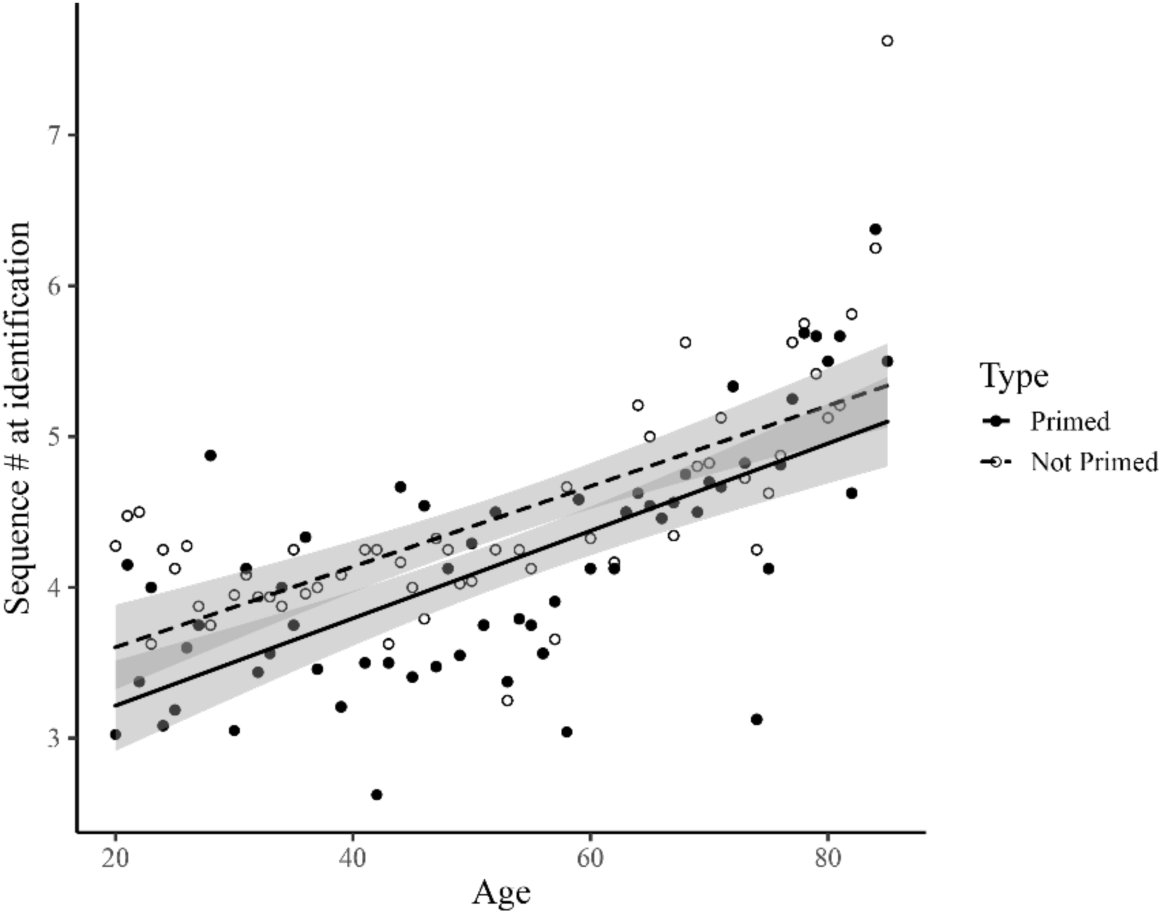
Performance in the Fragmented Identification Task. Participants needed to see more of the picture in order to correctly identify it as age increases (*p* < .001). However, they were consistently earlier at identifying a prime picture compared to a not primed picture (*p* < .001). There was therefore no effect of age on perceptual memory performance in this task (*p =* .744). Error clouds represents standard error.

Overall, there was no significant interaction between *Age* and *Primed*, suggesting that perceptual memory does not decline with age, consistent with the results reported for the WSC task.

### Conceptual Memory - Serial Reaction Time (SRT)

Following the methodology described in Neger and colleagues (2014), for each prime-target pair for each participant we calculated a facilitation score. The facilitation score is the reaction time to the prime picture divided by the reaction time to the target picture. This allows us to compare the prime-target reaction time ratio across participants without needing to correct for age-related slowing (Salthouse, 1996). Valid facilitation scores were restricted to those within 2.5 SD from the mean facilitation score within each participant. Figure 5 illustrates the change in facilitation score across the experimental session, averaged for each decade for illustrative purposes.

**Figure 5.**
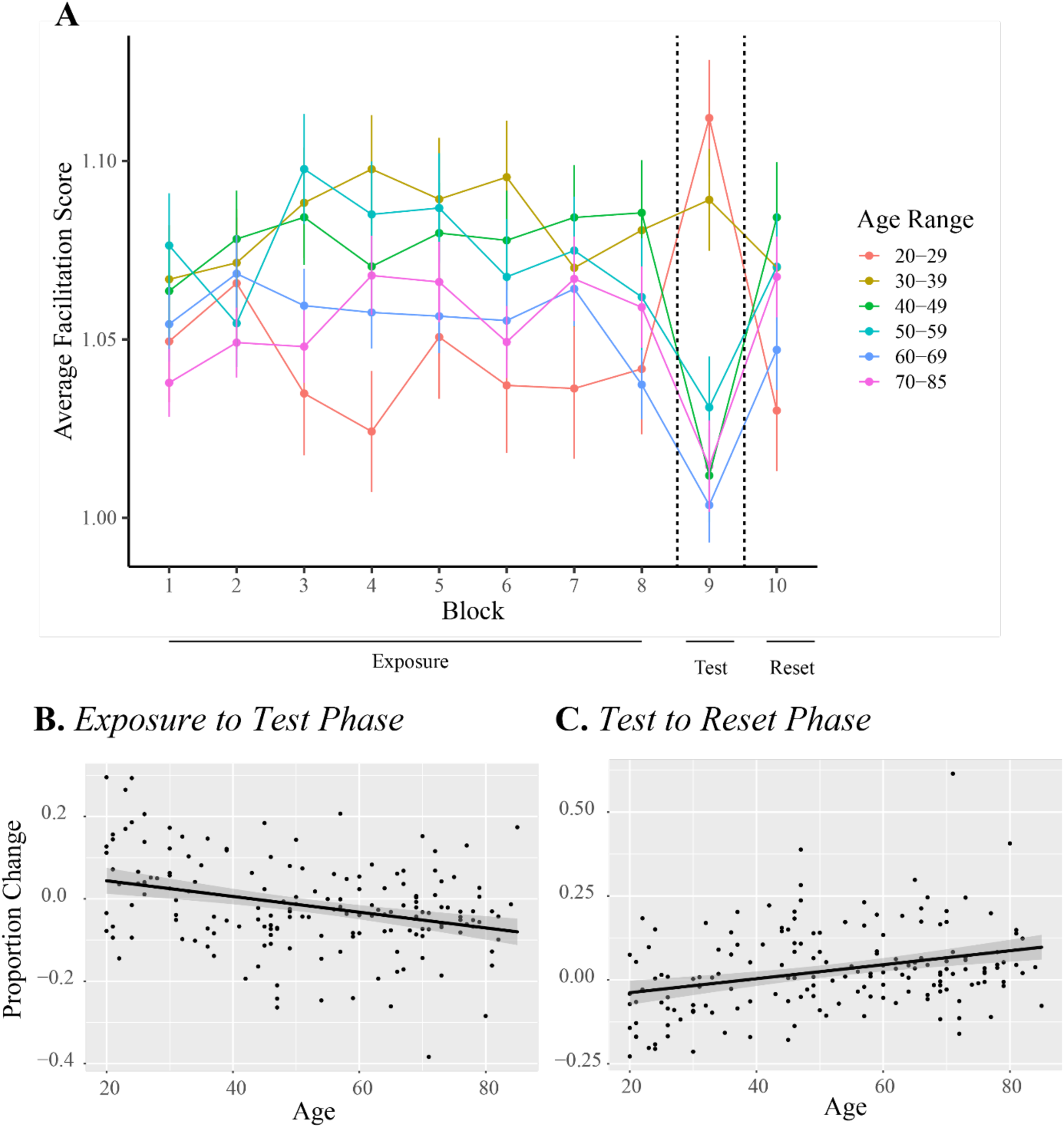
A. Average Facilitation Score for each block, averaged per decade for illustrative purposes. B. Percentage change in Facilitation Score for the Exposure to Test Phase (when the underlying pattern was changed) and C. for the Test to Reset Phase (when the underlying pattern was re-established). Both plots illustrate how percent change diverges from zero as age increases, suggesting differences in conceptual memory as a function of age.

Figure 5 shows how, throughout the course of the Exposure block, the facilitation score stabilizes as the participants get used to the task. For this reason, and in line with the methodology described in Neger and colleagues (2014), for analysis we only included blocks 7 and higher. We analyzed the data using linear mixed effects models, as each participant contributed multiple facilitation scores (one for each prime-target pair). This also allowed participants and items (spatial location) to be assessed as random factors to reduce the possibility of a Type I error (Barr et al., 2013). We started with a full model which included *Phase* (treatment coded with the Test block as reference group) in two-way interactions with *Age* and each of the three control measures, as well as *Age* in two-way interactions with each of the three control measures. Table 5 summarizes the results of the best model, which included an interaction of *Phase* and *Age* (as a linear factor), as well as an interaction of *Phase* with *Working Memory*. The model included participant and item as random intercepts, but the model did not converge when random slopes were included.

**Table 5.**
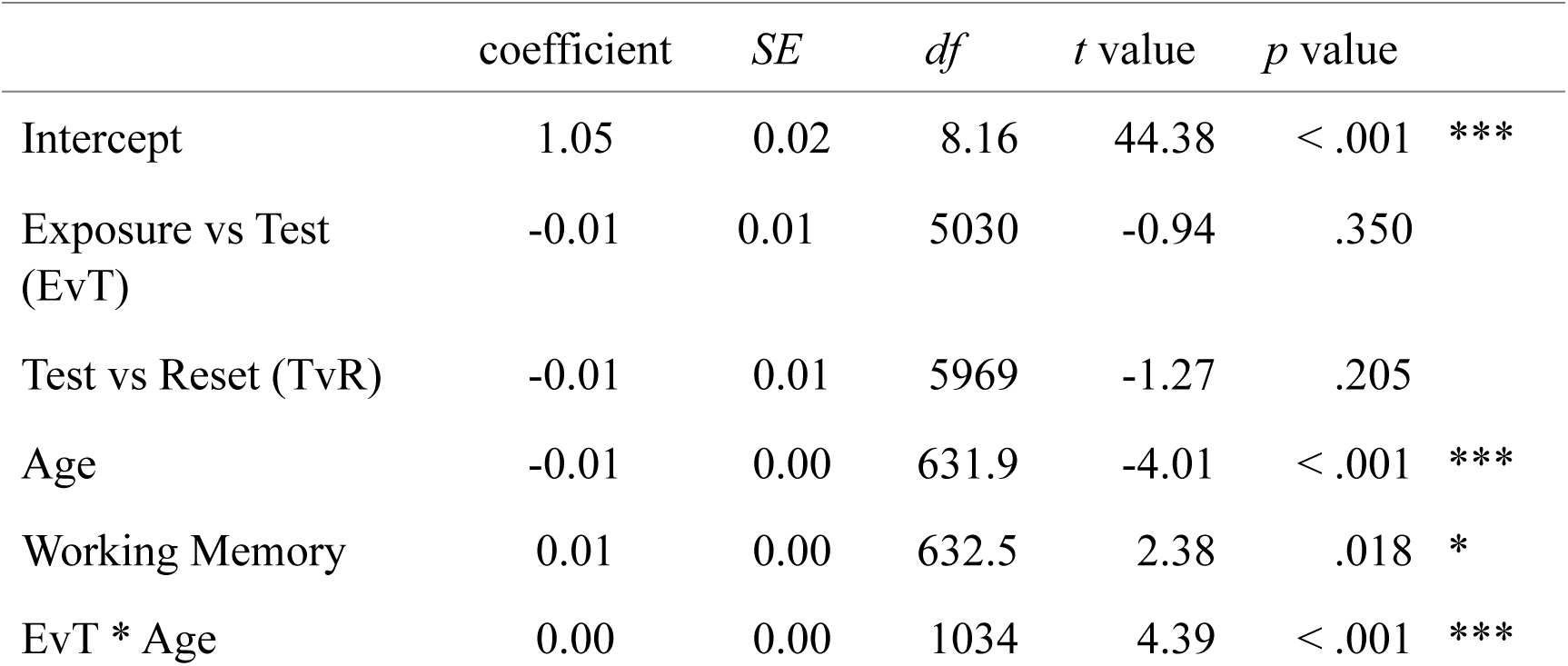
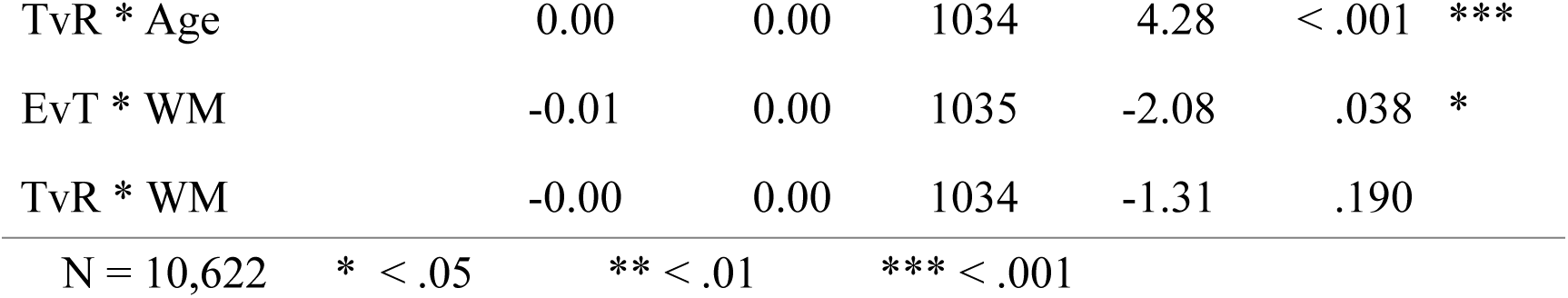
Summary of the best linear model for the Serial Reaction Time Task for Facilitation Score of the Exposure (blocks 7 – 8), Test (blocks 9 – 10), and Reset Phase (blocks 11 – 12).

|  | coefficient | SE | df | t value | p value |  |
| --- | --- | --- | --- | --- | --- | --- |
| Intercept | 1.05 | 0.02 | 8.16 | 44.38 | < .001 | *** |
| Exposure vs Test (EvT) | -0.01 | 0.01 | 5030 | -0.94 | .350 |  |
| Test vs Reset (TvR) | -0.01 | 0.01 | 5969 | -1.27 | .205 |  |
| Age | -0.01 | 0.00 | 631.9 | -4.01 | < .001 | *** |
| Working Memory | 0.01 | 0.00 | 632.5 | 2.38 | .018 | * |
| EvT * Age | 0.00 | 0.00 | 1034 | 4.39 | < .001 | *** |
| TvR * Age | 0.00 | 0.00 | 1034 | 4.28 | < .001 | *** |
| EvT * WM | -0.01 | 0.00 | 1035 | -2.08 | .038 | * |
| TvR * WM | -0.00 | 0.00 | 1034 | -1.31 | .190 |  |
| N = 10,622 | * < .05 | ** < .01 | *** < .001 |  |  |  |

Conceptual memory was defined as a drop in facilitation score in the Test Phase (blocks 9 – 10) compared to the facilitation score at the end of the Exposure Phase (blocks 7 –8). This is because the underlying pattern in the test phase is different compared to the exposure phase, and hence the participant’s reaction time to the target location should be slower. If this were different depending on age, the model would represent that as a significant interaction between *Exposure versus Test Phase* and *Age*, which is indeed what we find (β = 0.00, *p* < .001). Figure 5B illustrates this as a percent change between the participant’s average facilitation score between the Exposure Phase and Test Phase. The figure shows that as participants get older, their drop becomes more defined, suggesting they become more vulnerable to the change in the underlying pattern.

We also included the comparison between the Test Phase and the Reset Phase (blocks 11 – 12). In the Reset Phase, the same underlying pattern as that in the Exposure Phase is re-introduced, and hence an increase in facilitation score is expected. This is a control measure to show that the participants did indeed learn the underlying pattern during the Exposure Phase. The model again shows a significant interaction of this effect with *Age* (β = 0.00, *p* < .001). Figure 5C shows that as the participants increase in age, their facilitation score increases more. This suggests that the older participants did not attempt to change their predictions during the Test Phase to account for the new change in pattern. It is logical that the younger participants were able to pick up on this change, and learn this change, faster and hence showed an overall smaller change in their facilitation scores.

Overall, we show a significant effect of age on the performance in the SRT task.

### Structural Priming Task

We define short-term priming as the influence that processing a prime has on the syntactic choice of the immediately following target, and long-term priming as the influence all past produced syntax has on the syntactic choice of the current target. Long-term priming was therefore calculated as the proportion of passives out of the total transitive responses produced on the target trials before the current target trial. A positive and significant *Long-term Priming Magnitude* therefore suggests that the proportion of passives previously produced positively influences the probability of producing a passive on the current target trial. Previous studies have also referred to this as *Cumulative Passive Proportion* (Heyselaar et al., 2015; Jaeger & Snider, 2008). We did not calculate this for active primes as very few previous priming studies have shown a significant active priming effect, and hence we did not consider this variable to model learning.

Previous studies have shown a non-linear correlation between the syntactic choice on the current target sentence and past produced syntax (Heyselaar et al., 2015, 2017; Jaeger & Snider, 2008, 2013), we therefore decided to analyse the data using generalised additive mixed models (GAMM). Unlike ANOVAs or generalized mixed-effects regression (GLMER), GAMM does not assume linearity (although it can find a linear form if supported by the data). Instead, GAMM strikes a balance between model fit and the smoothness of the curve using either error-based or likelihood-based methods in order to avoid over-or under-fitting. Thus, the data guide the functional form. The *p*-value provided therefore indicates whether or not the curve is significantly different from zero. Additionally, GAMM also allows the inclusion of random effects to capture the dependencies between repeated measures.

The full model contained the two-way interaction *Prime* (treatment-coded, baseline trials as reference group) by *Age* in addition to interactions of *Prime* with each of the three control measures. *Prime* represents the active, passive, or baseline structure the participant produced during the prime trials and whether this affects the structure produced on the subsequent target trials. This therefore measures the short-term priming effect.

The best model included main effects of *Prime, Age,* and two-way interactions between *Age* and *Long-term Priming Magnitude* and *Age* and *Prime. Prime* was modelled as a linear predictor as the estimated degrees of freedom were 1. The remaining predictors were modelled with smoothers, using the default underlying base functions (thin plate regression for *Prime* by *Age* and cubic regression for *Long-term Priming* by *Age*). We included per-participant and per-item random smooths. The function gam.check was used to ensure adequate k was used for each predictor.

Table 6 illustrates the results from the best model but including the *Prime* by *Age* interaction for illustrative purposes.

**Table 6.** Summary of the best generalised additive mixed effects model for the structural priming task. LtPM - Long-term Priming Magnitude

|  | Estimate | SE | z value | p value |  |
| --- | --- | --- | --- | --- | --- |
| Intercept | -4.55 | 0.18 | -25.39 | < .001 | *** |
| Active Prime (AP) | -0.06 | 0.15 | -0.39 | .699 |  |
| Passive Prime (PP) | 0.47 | 0.14 | 3.31 | < .001 | *** |
| Age | 0.11 | 0.06 | 1.94 | .052 |  |

| B. Smooth Terms |  |  |  |  |  |
| --- | --- | --- | --- | --- | --- |
|  | edf | Ref.<br>df | X<sup>2</sup><br>value | *p* value |  |
| Smooth for Age - Baseline | 2.03 | 2.50 | 3.02 | .390 |  |
| Smooth for Age - AP | 0.00 | 0.00 | 0.00 | .994 |  |
| Smooth for Age - PP | 2.94 | 3.59 | 4.36 | .258 |  |
| Smooth for Age - LtPM | 10.89 | 12.4 | 452.90 | < .001 | *** |
| Random effect for<br>Participants | 31.81 | 152.0<br>0 | 50.19 | < .001 | * |
| Random effect for Items | 32.62 | 56.00 | 88.70 | < .001 | *** |
| N = 8587 * < .05 ** < .01 *** < .001 |  |  |  |  |  |

The negative estimate for the intercept indicates that in the baseline condition (intransitive prime followed by transitive target) active responses were more frequent than passive responses. Following passive primes, more passive responses were produced compared to baseline (*p* < .001). Following active primes, there was no increase in active responses compared to baseline (*p* = .699). This is the standard pattern of results reported in the literature (e.g., Bernolet et al., 2016; Bock, 1986; Ferreira & Bock, 2006). There was no interaction of *Prime* with *Age* (*p* > .258; Figure 6A) suggesting that the short-term priming effect does not vary as a function of age.

**Figure 6.**
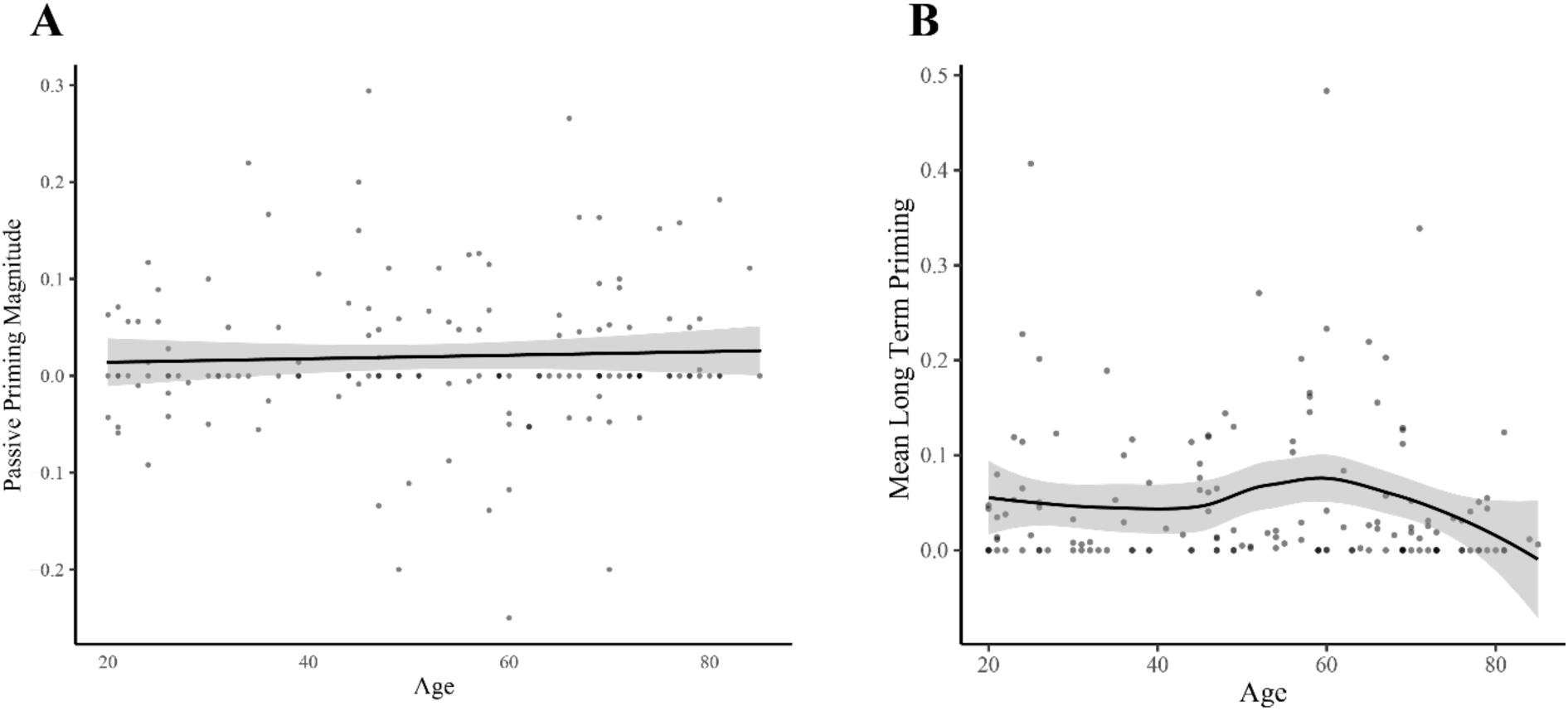
Performance in the Structural Priming Task A. Short-term priming. We observed no significant effect of age on short-term priming magnitude (p > .105). B. Long-term priming. We observe a non-linear influence of age on long-term priming magnitude between 50 and 70 years. Error clouds/bars represent standard error.

We do observe a significant interaction between *Long-term Priming Magnitude* and *Age*, suggesting that there is an effect of age on long-term priming (Figure 6B). The model and figure suggest that the interaction between *Age* and *Long-term Priming Magnitude* is not a simple quadratic correlation; however, the interaction is significant (*p < .*001) suggesting there is an effect of age on long-term priming. The figure suggests that after 40 years of age, there is an increase in the average long-term priming magnitude; which means that past use of a passive structure plays a stronger role in predicting whether the current utterance will use a passive structure. The figure suggests that after age 60, this effect disappears, although it is hard to conclude why this happens from our data.

Overall, we see no effect of age on short-term priming, while we see a clear effect of age on long-term priming.

### Interaction of Conceptual and Perceptual Memory on Syntactic Priming

As a final element to support our hypothesis that perceptual memory underlies short-term priming and that conceptual memory underlies long-term priming, we attempted to correlate the individual participant scores from each memory task with their short-term and long-term priming performance.

In order to combine the scores, we conducted a Principal Components Analysis (PCA) on the datasets used to model the word-stem completion (WSC), the fragmented identification, and the serial reaction time (SRT) tasks, respectively. For the WSC, we calculated the *difference* in reaction time between the Primed and Unprimed conditions, for the fragmented identification task, we calculated the *difference* in the number of pictures seen between the primed and unprimed conditions, and for the SRT, we calculated the *difference* in the facilitation score between the Exposure and Test phase, as well as the Test and the Reset phase. We ensured that all the data used was the same as the data entered into the model for each respective task. In total, our dataset included four variables: WSC difference, fragmented identification difference, Exposure-Test difference, and Test-Reset difference. As certain participants were removed in individual tasks due to outliers, we only included the data from 158 participants that had scores in all four variables.

A PCA was conducted on the 4 items with orthogonal rotation (varimax). Bartlett’s test of sphericity (χ^2^ (158) = 211.36, *p* < .001) indicated that correlations between the items were sufficiently large for PCA. Both the Kaiser rule (eigenvalues > 1.0) and parallel analysis (a simulation procedure using randomly generated data to estimate the number of components), suggested a 2 factor model. Table 7 reports the factor loadings for each of the 2 components, as well as the proportion of variance and Cronbach’s α. Note that the Cronbach’s αfor the perceptual memory component is extremely low. However, extracting three components (and therefore separating WSC and the fragmented identification task) provided the same final conclusion and hence here we will report the results that match our hypothesis.

**Table 7.** Factor loadings of the SRT, WSC and Fragmented Identification task scores. No loadings are omitted (N = 158). EvT – Exposure vs Test Phase (SRT Task); TvR – Test vs Reset Phase (SRT Task)

| Item | *Varimax rotated factor loadings* |  |
| --- | --- | --- |
|  | Conceptual Memory | Perceptual Memory |
| TvR Facilitation Score Diff. | **0.96** |  |
| EvT Facilitation Score Diff. | **0.95** |  |
| WSC RT Diff. |  | **0.74** |
| FI PicturesSeen Diff |  | **0.69** |
| Eigenvalues | 1.88 | 1.03 |
| Proportion variance | 0.47 | 0.26 |
| $\alpha$ | .92 | .01 |

The factor scores were then added to the structural priming data set, which resulted in a dataset for 138 participants who had scores in all variables. We created a hypothesis-driven model, such that the model included a main effect for *Prime,* the *Conceptual* and *Perceptual Memory* components, as well as two-way interactions between *Prime* and *Perceptual Memory*, and *Long-term Priming Magnitude* and *Conceptual Memory. Prime* and its interaction was, again, modelled as a linear predictor, whilst *Long-term priming* and its interaction were modelled as smooth terms. Table 8 reports the results of this model.

**Table 8.**
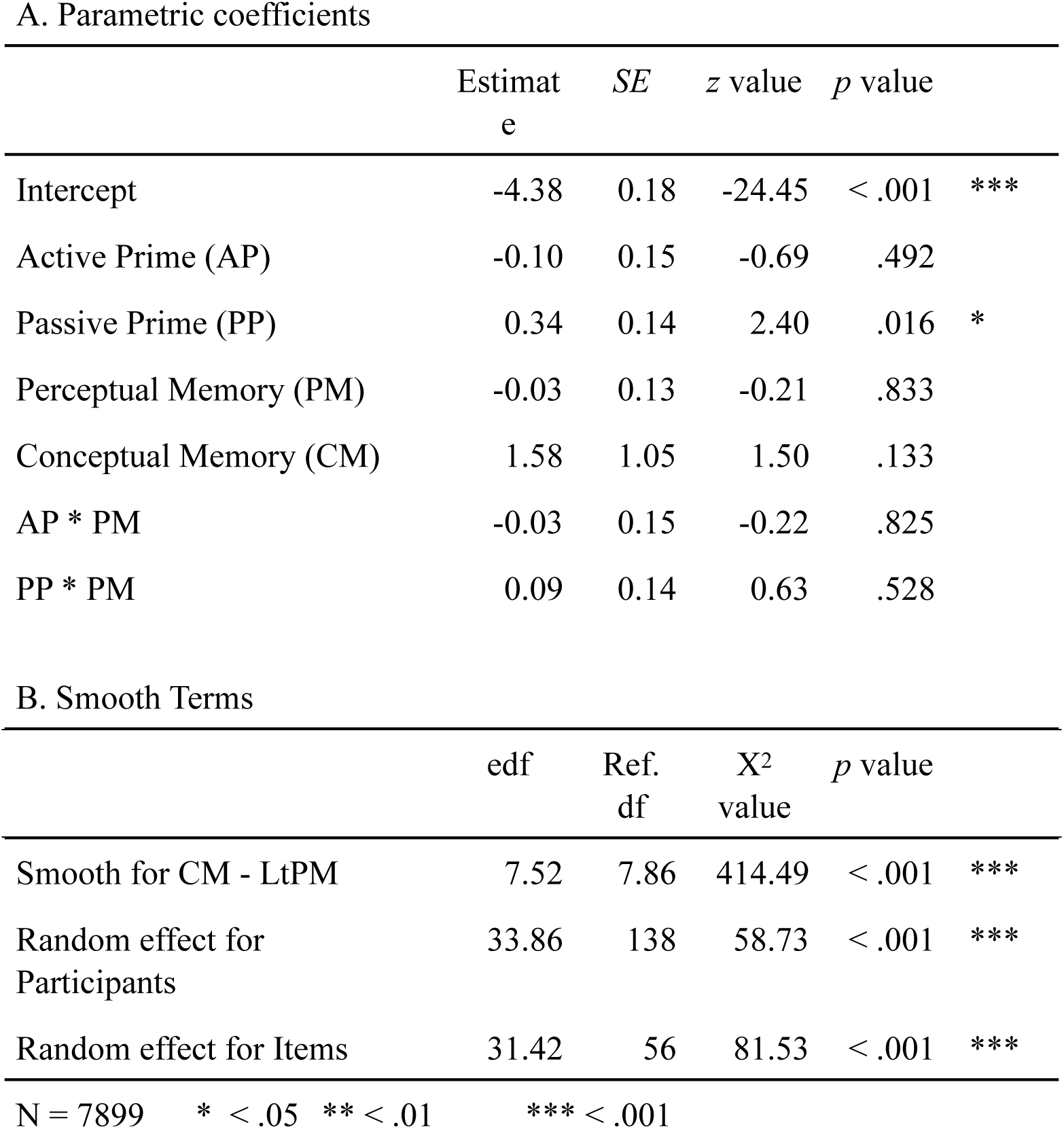
Summary of the generalised additive mixed effects model for relationship between the structural priming task and the non-declarative memory components. LtPM - Long-term Priming Magnitude

A. Parametric coefficients
|  | Estimate | SE | z value | p value |  |
| --- | --- | --- | --- | --- | --- |
| Intercept | -4.38 | 0.18 | -24.45 | < .001 | *** |
| Active Prime (AP) | -0.10 | 0.15 | -0.69 | .492 |  |
| Passive Prime (PP) | 0.34 | 0.14 | 2.40 | .016 | * |
| Perceptual Memory (PM) | -0.03 | 0.13 | -0.21 | .833 |  |
| Conceptual Memory (CM) | 1.58 | 1.05 | 1.50 | .133 |  |
| AP * PM | -0.03 | 0.15 | -0.22 | .825 |  |
| PP * PM | 0.09 | 0.14 | 0.63 | .528 |  |

B. Smooth Terms
|  | edf | Ref. df | X<sup>2</sup> value | p value |  |
| --- | --- | --- | --- | --- | --- |
| Smooth for CM - LtPM | 7.52 | 7.86 | 414.49 | < .001 | *** |
| Random effect for Participants | 33.86 | 138 | 58.73 | < .001 | *** |
| Random effect for Items | 31.42 | 56 | 81.53 | < .001 | *** |
| N = 7899 * < .05 ** < .01 *** < .001 |  |  |  |  |  |

The model shows no significant interaction between short-term priming and perceptual memory (*p* = .528), suggesting that the participant’s perceptual memory performance does not influence their likelihood of producing a passive target after having produced a passive prime. However, there is a significant interaction between long-term priming and conceptual memory (*p* < .001), suggesting that performance in the serial reaction time task is correlated to the participant’s long-term priming magnitude.

## Discussion

In this study we aimed to test the proposal that both long-term and short-term structural priming are supported by non-declarative memory. Specifically, we tested our hypothesis that long-term priming is supported by conceptual memory whereas short-term priming is supported by perceptual memory - both subcomponents of the non-declarative memory system. We investigated how the magnitude of these two priming effects varied with the age of the participants. Previous studies in the memory literature have suggested that the two subcomponents of non-declarative memory age differently by demonstrating age-related decline in conceptual memory but not in perceptual memory. We therefore investigated how age-related decline in these two components of non-declarative memory relates to performance in structural priming, in order to elucidate how non-declarative memory supports two seemingly different components of structural priming.

Our tasks and their key findings are summarized in Table 9. The results of this study show that there is no age-related effect in either the perceptual memory tasks or the short-term structural priming magnitude. In contrast, our results do demonstrate an age-related effect on the performance in both the conceptual memory tasks and the long-term priming magnitude. This pattern of results is consistent with our hypothesis, which locates short-term structural priming in perceptual memory and long-term structural priming in conceptual memory. We elaborate on this below.

**Table 9.** Summary of key results for the non-declarative memory tasks

| **Task** | **Non-declarative Memory Component Measured** | **Key Results** |
| --- | --- | --- |
| Word-stem completion | Perceptual memory | No age effect |
| Fragmented Identification | Perceptual memory | No age effect |
| Serial Reaction Time | Conceptual memory | More pronounced changes in performance with increasing age |
| Structural Priming | Short-term: Perceptual memory | No age effect |
|  | Long-term: Conceptual memory | Increased long-term priming magnitude between 50 – 70 years |

Firstly, our study shows a clear difference in the age-related effects on performance in the two subcomponents of non-declarative memory, with changes in performance in conceptual memory tasks but not in perceptual memory tasks. In the SRT task, we observed more pronounced changes in the facilitation score of the older participants compared to their younger peers. This change does not necessarily indicate that older participants have a better conceptual memory than younger participants, but that they are slower to adapt to new patterns. During the SRT task, they did not adapt their predictions even when it was clear there was a new underlying pattern, and hence the percent decrease from the Exposure to Test phase (3.59%, SD: 10.48%) is close to the percent increase from Test to Reset phase (6.39%, SD: 12.55%) as the older participants did not update their predictions.

Our results contribute substantially to the broader question of age-related changes in non-declarative memory. As discussed in the Introduction, previous literature has provided equivocal conclusions whether either non-declarative memory decreases with age. A major factor was that previous studies would directly compare younger participant groups to older groups, with the average age of the older participants varying from early 60’s (Howard, Heisey, & Shaw, 1986; Neger, Rietveld, & Janse, 2014; Schugens, Daum, Spindler, & Birbaumer, 2007) to late 80s (Davidson, Zacks, & Ferreira, 2003; Davis et al., 1990; Karlsson, Adolfsson, Börjesson, & Nilsson, 2003; Light, Kennison, & Healy, 2002; Light, La Voie, Valencia-Laver, Albertson Owens, & Mead, 1992). In our study, we therefore opted for a longitudinal design, and tested participants between 20 - 85 years. Our results show that there is a significant age-related decline, and that this already starts around 50 years of age. Additionally, we find an opposite trend to that found in Maki and colleagues (1999), one of the only empirical studies that do look at age as a continuous variable in their study design. They observed a significant decrease in age for the perceptual memory task (Fragmented Object Identification) and no change for their conceptual memory task (Category Exemplar). This could be due to difference in task design and execution (see La Voie & Light, 1994 for a discussion on this for the word-stem completion task), and therefore more empirical work is necessary to further standardise non-declarative memory tasks and report their results on age-related decline.

Second, our findings demonstrate an age-related effect on performance in long-term structural priming but not in short-term structural priming, suggesting a link between the temporal characteristics of structural priming and the two subcomponents of non-declarative memory. In our data, the older participants were more likely to produce a passive on the current trial if they had produced passives previously. This again hints that these participants are less able to adapt to changes and instead are more likely to repeat what they have done in the past, similar to the SRT task. This interpretation is purely *post hoc* given the data but there are already numerous studies arguing for long term priming to be supported by implicit learning (Chang, Dell, & Bock, 2006; Chang, Janciauskas, & Fitz, 2012; Jaeger & Snider, 2013; Reitter, Keller, & Moore, 2011, inter alia).

Finally, we used principal components analysis to support the existence of a link between the performance in the conceptual and perceptual memory tasks and short- and long-term priming magnitude. Our results showed a significant interaction between long-term priming and the conceptual memory component (*p* < .001), however, we observed no correlation between short-term priming and the perceptual memory component. One explanation is that there were problems in having the two perceptual memory components load onto the same factor (Cronbach’s α: .01), even though both tests are commonly used to measure perceptual memory. Therefore, it could be that the factor did not capture perceptual memory as we intended, and as such the existence or lack of a correlation with the short-term memory magnitude should be interpreted with caution. However, the lack of an age-related effect in performance for the perceptual memory tasks as well as in the short-term priming magnitude does suggest that the underlying memory systems could be related. A replication is therefore necessary before any strong conclusions can be drawn about the link between perceptual memory and short-term syntactic priming.

Our study is one of the first to show a clear dissociation in how increasing age affects short-term and long-term priming, as even though the short-term priming magnitude is lower than other structural priming studies, we are still about to see a clear age effect for the long-term compared to the short-term priming magnitudes. Many developmental studies have shown that children show higher priming magnitude and a higher tendency to prime compared to adult studies (Branigan & Messenger, 2016; Kidd, 2012), and thus priming ability already declines from 3 – 4 year olds to the student population. We have provided insight into how this decline continues as we age.

Our study is an important first step towards providing evidence of a connection between non-declarative memory and both temporal characteristics of structural priming. Future studies will focus on the nature of the link between structural priming and different memory components. In our study we focused on abstract priming in transitives; in order to establish our proposed non-declarative memory account, the findings reported here should be replicated with other syntactic structures (i.e. datives), as well as lexical overlap reintroduced into the paradigm to determine what role it plays in supporting structural priming ability as we age. Additionally, as older adults are known to recruit additional brain areas to compensate for lost efficiency elsewhere (Reuter-Lorenz & Park, 2014; Wingfield, Peelle, & Grossman, 2003), neuroimaging studies will shed additional light on if/how the brain networks underlying structural priming differ as we get older.

Previous studies using patients with amnesia have highlighted the supporting role that the non-declarative memory system plays in structural priming. Both Ferreira and colleagues (2008) and Heyselaar and colleagues (2017) have demonstrated a robust priming magnitude when testing amnesia patients on either double-object/prepositional-object or active/passive structural priming tasks. Therefore, it has been accepted that structural priming is supported by non-declarative memory. However, as structural priming itself is made up of both a short-term and a long-term component, models have been struggling to explain how one system could support both of these temporally distinct characteristics. We suggest these two different structural priming components are subserved by different non-declarative memory components, which has important implications for theoretical accounts of structural priming.

Our results suggest that perceptual memory underlies the short-term component and that conceptual memory underlies the long-term component of structural priming. This is supported by a combination of different existing structural priming models: We propose a residual activation account, based in non-declarative memory (similar to that proposed by Pickering and Branigan, 1998; Malhotra, 2009; Reitter et al., 2001), for short-term priming and a non-declarative learning account (similar to that proposed by Chang et al., 2006; Jaeger & Snider, 2013) for long-term priming. The information transfer between these components is based on the information transfer proposed above in the Reitter model. They model priming as spreading activation, and assume that lexical forms persist in a working memory buffer in order to process their semantic contributions, e.g., for the duration of a sentential unit, until they are replaced by other lexical forms. Similarly, they propose that semantic information can persist even beyond the utterance. By virtue of being in a buffer, lexical forms and semantic information can then spread activation from the buffer to associated chunks in memory, such as syntactic categories. The more frequent the syntactic category is, the greater its prior probability. Non-declarative memory works in a similar fashion: Perceptual memory measures the residual activation of a previously processed item that persists, represented as the decreased reaction time when this item is processed a second time. With repeated exposures to this item, however, a link can be made to conceptual memory, if there are underlying links between the items (Poldrack, Selco, Field, & Cohen, 1999). Therefore, repeated exposures to the same item(s) enhances this link, influencing its baseline-activation and hence the probability of it influencing an upcoming response, mostly measured as a decrease in processing latency of the item or construct (as in the serial reaction time task, for example). This type of model also supports structural priming effects seen in reaction times (Corley & Scheepers, 2002; Segaert, Weber, Cladder-Micus, & Hagoort, 2014; Segaert et al., 2016; Segaert et al., 2011; Wheeldon & Smith, 2003), an important and robust phenomenon usually not included in models of structural priming. Segaert and colleagues (2011, 2014, 2016) have proposed a two-stage competition model to explain the reaction time effects, the basis of which is very similar to Reitter and colleagues (and our) proposal of a base-level residual activation that spreads and is updated depending on repeated exposures. Of course, Reitter and colleagues propose explicit memory influences in their baseline-activation, whereas we propose the whole system be fully based in non-declarative memory.

The observed influences of age on structural priming can also be explained with the same non-declarative mechanisms as those we described above. There are two aging theories that speak to the tasks we used in this study, the processing-speed theory (Salthouse, 1996) and the transmission deficit hypothesis (Mackay & Burke, 1990). The processing-speed theory (also referred to as general slowing), proposes that information from different sources may become available to a central processor so slowly that the earlier information has decayed or is no longer active by the time the later information arrives. In terms of structural priming, this could suggest that the residual activation of a structure is not available long enough to influence updating the statistical knowledge of that structure, and hence its baseline-activation is never changed. Therefore, we would see the short-term priming effect, but a diminished long-term priming effect. The transmission-deficit hypothesis is very similar in this regard: The authors suggest that the encoding of new memories and retrieval of existing memories depends on the rate of transmission across the connections linking representational units in memory. They provide a priming related example in their text (Burke & Mackay, 1997; p. 1852): “Priming is a form of subthreshold excitation that prepares a unit for activation or retrieval, and the rate of priming transmission depends on the strength of connections among units. Aging is postulated to weaken connection strength.” Again, the weakening of connection strength also explains why we see robust short-term priming effects, but weak long-term priming effects for the oldest age groups.

In conclusion, our study supports our proposal that non-declarative memory underlies two distinct structural priming effects: Short-term and long-term priming. The perceptual component of the non-declarative memory system supports short-term priming effects, whereas the conceptual component supports long-term priming effects. In this study, we focused solely on abstract priming (no lexical overlap between prime and target trials); it remains to be investigated how these mechanisms may change with lexical overlap. Our study is also the first to show divergent effects of age on the two components of structural priming, an important characteristic that needs to be included in models of structural priming. It therefore provides important new insights into the relationship between non-declarative memory and language production and, in addition, new insights into how both memory and language are affected by age.

## Acknowledgments

We would like to thank Denise Clissett, the coordinator of Patient and Lifespan Cognition participant database at the University of Birmingham, for recruiting and scheduling participants. We would also like to thank Emma Sutton, Marissa McCallum, Bessie McDonald-Phelps, Emily Robinson, Ellie Cooper, and Charlotte Poulisse for their help with collecting the data. We thank Florian Jaeger for his advice on a previous version of this manuscript.

